# *DEFECTIVELY ORGANIZED TRIBUTARIES 5* is not required for leaf venation patterning in *Arabidopsis thaliana*

**DOI:** 10.1101/2022.07.19.500632

**Authors:** Daniela Vlad, Jane A. Langdale

**Affiliations:** Department of Plant Sciences, University of Oxford, South Parks Rd, Oxford OX1 3RB, UK

**Keywords:** Leaf development, gene editing, venation patterning, *Arabidopsis thaliana*

## Abstract

The search for genetic regulators of leaf venation patterning started over thirty years ago, primarily focussed on mutant screens in the eudicotyledon *Arabidopsis thaliana*. Developmental perturbations in either cotyledons or true leaves led to the identification of transcription factors required to elaborate the characteristic reticulated vein network. An ortholog of one of these, the C2H2 Zn finger protein DEFECTIVELY ORGANIZED TRIBUTARIES 5 (AtDOT5), was recently identified through transcriptomics as a candidate regulator of parallel venation in maize leaves. To elucidate how *AtDOT5* orthologs regulate vein patterning, we generated three independent loss of function mutations by gene editing in Arabidopsis. Surprisingly, none of them exhibited any obvious phenotypic perturbations. To reconcile our findings with earlier reports, we re-evaluated the original *Atdot5-1* and *Atdot5-2* alleles. By genome sequencing, we show that reported mutations at the *Atdot5-1* locus are actually polymorphisms between Landsberg *erecta* and Columbia ecotypes, and that other mutations present in the background most likely cause the pleiotropic mutant phenotype observed. We further show that a T-DNA insertion in the *Atdot5-2* locus has no impact on leaf venation patterns when segregated from other T-DNA insertions present in the original line. We thus conclude that *AtDOT5* plays no role in leaf venation patterning in Arabidopsis.

**Significance statement:** An understanding of gene function is often derived on the basis of loss of function mutant phenotypes and thus correct identification of mutated loci is crucial. Through gene editing we reveal previous mis-identification of a causative mutation that led to inappropriate functional assignation for the *DEFECTIVELY ORGANIZED TRIBUTARIES 5 gene* in *Arabidopsis thaliana*.

## INTRODUCTION

Vascular plants rely on a complex network of veins to transport water, nutrients and sugar throughout the plant and to serve as structural support. How the vascular network is established and maintained has been the focus of research for decades, spanning disciplines from physiology and development to hydraulics and mathematics (reviewed in Nelson and Dengler, 1997, Sack and Scoffoni, 2013, De Rybel *et al*., 2016, Perico *et al*., 2022). Early physiological studies revealed that auxin induces vascular formation after wounding, leading to the hypothesis that canalization of auxin flow through developing veins facilitates self-organization of the vascular network (Sachs, 1969, Sachs, 1981). Attempts to validate this hypothesis at the molecular level are still ongoing, with new discoveries alternately supporting or refuting the possibility (reviewed in Rolland-Lagan and Prusinkiewicz, 2005, Bennett *et al*., 2014, Ravichandran *et al*., 2020, Perico *et al*., 2022). Central to these endeavours are screens for mutants with perturbed vascular patterning that enable causative mutations to be identified and thus genetic regulators of the patterning process to be revealed. Mutants with perturbed vascular patterning in roots and/or shoots have been reported in both eudicotyledons and monocotyledons, with most studies to date focussed on the eudicot species *Arabidopsis thaliana* (e.g. Hardtke and Berleth, 1998, Candela *et al*., 1999, Carland *et al*., 1999, Scarpella and Meijer, 2004, Petricka *et al*., 2008, Smillie *et al*., 2012).

The extent to which patterning mechanisms are conserved in eudicot leaves that elaborate reticulate vein networks and monocot leaves that develop parallel veins is unknown, however, transcriptomic studies identified an ortholog of the Arabidopsis *DEFECTIVELY ORGANIZED TRIBUTARIES 5* (*AtDOT5*) gene as a candidate regulator of leaf venation patterning in maize (Wang *et al*., 2013). This observation suggests that aspects of the patterning process may be shared in eudicots and monocots. The Arabidopsis *defectively organized tributaries* (*Atdot*) mutants were isolated from an extensive forward genetics screen for altered vein patterning in young leaves of ∼30000 individuals (Petricka *et al*., 2008). The plants were derived from three different mutagenized populations, two generated in the Columbia-0 (Col-0) background using either diepoxybutane (Clay and Nelson, 2005) or activation tagging-mutagenesis (Weigel *et al*., 2000), and one obtained by *Dissociation* (*Ds*) transposon-mutagenesis in the Landsberg *erecta* (L*er*) background (Bancroft *et al*., 1992). The pleiotropic *Atdot5-1* mutant was isolated from the *Ds* mutagenized *Ler* background and was classically mapped to the At1g13290 (*AtDOT5*) locus (Petricka *et al*., 2008), which encodes for a WIP family C2H2 Zn finger protein with predicted transcription factor activity (Appelhagen *et al*., 2010). *Atdot5-1* mutants exhibited narrower leaves with misaligned veins, delayed leaf initiation, reduced apical dominance, short roots and enhanced auxin sensitivity. The *Atdot5-1* allele differed from wild-type at four amino acid positions, all outside of the C2H2 and WIP domains. A second allele *Atdot5-2*, contained a T-DNA insertion upstream of the C2H2 domain that conditioned an embryo lethal phenotype (Alonso *et al*., 2003, Petricka *et al*., 2008). A second T-DNA insertion in the line containing the *Atdot5-2* allele was subsequently reported in the 5’UTR of At2g26740 (https://abrc.osu.edu/stocks/631530) which encodes one of the Arabidopsis Soluble EPOXIDE HYDROLASES (AtSEH) (Kiyosue *et al*., 1994, Pineau *et al*., 2017). Constitutive expression of the *AtDOT5* genomic sequence from wild-type L*er* was reported to only partially complement the leaf initiation defects of the *Atdot5-1* mutant (Petricka *et al*., 2008). To date, the mechanism by which AtDOT5 regulates vein patterning in Arabidopsis has not been elucidated.

Given the potential for conserved function of *AtDOT5* orthologs in both eudicot and monocot leaf vein patterning, we sought to determine the mechanism of gene function. To this end we used CRISPR/CAS9 to generate three independent loss of function alleles in Arabidopsis. Here we report the characterization of those mutants and a re-analysis of the original mutant alleles.

## RESULTS & DISCUSSION

### *Gene edited loss of function* Atdot5 mutants *show no vein patterning or morphological defects*

To generate null alleles of *AtDOT5*, CRISPR/Cas9 was used in conjunction with a single guide RNA that was designed to target the first exon of the At1g13290 locus. The guide was predicted to bind 100 bp downstream from the ATG and to induce mutations that would disrupt both the WIP and the C2H2 zinc finger domains (Figure 1A). T1 plants were screened for potential mutations by genomic PCR and three different loss of function alleles were identified in which deletions or insertions led to a premature stop codon (Figure 1A). Two alleles, a 5 bp deletion (C11.1_4) and a 1 bp insertion (C11.4_A3), arose in the same background (line C11). In order to segregate away any potential interactors from the shared C11 background that might influence the mutant phenotype, the two lines were independently backcrossed as the male parent to wild-type Col-0 and then selfed to F2. The third allele, a 1 bp deletion, was isolated as a heterozygous, transgene free T2 line and was fixed as homozygous (F12.12_5) in the T3 generation. For all three mutant alleles, segregating wild-type siblings were fixed in the same generation for use as controls in phenotyping experiments.

**Figure 1.**
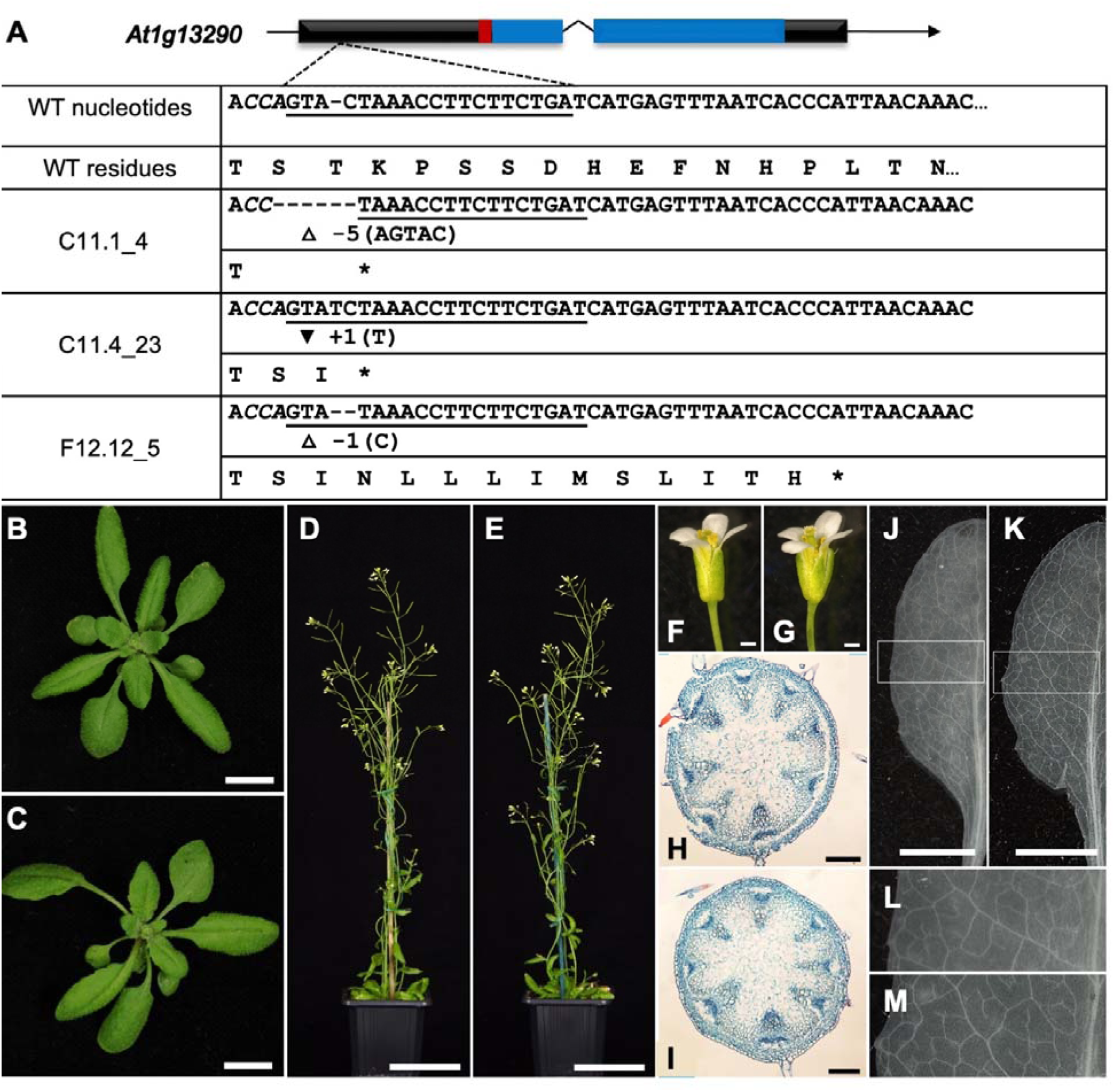
CRISPR generated *Atdot5* loss of function mutants resemble wild type plants. **A)** Schematic of the *AtDOT5* locus targeted for mutagenesis and representation of three edited alleles. Black boxes depict exons with the overlying red box depicting the WIP domain and the blue boxes depicting the zinc finger domains. The sgRNA complementary sequence is underlined and the PAM site is shown in bold italic. White triangles mark deletions and the black inverted triangle marks an insertion. **B-M)** Phenotypic characterization of segregating wild-type (F12.12_3) (B, D, F, H, J & L) and homozygous *Atdot5* loss of function mutants (F12.12_5) (C, E, G, I, K & M). Plants were imaged prior to bolting (B, C) and after flowering (D, E). Flower morphology appeared normal (F, G) and vascular patterning in both stems (H, I) and leaves (J-M) was similar in wild-type and loss of function lines. L & M are insets of J & K respectively, as indicated. Scale bars: 1 cm (B, C); 5 cm (D, E); 0.5 mm (F, G); 200μm (H, I) and 5 mm (J, K).

To characterize the mutant phenotype, homozygous mutants from all three independent lines were grown to maturity alongside wild-type controls. Unexpectedly, mutant plants were developmentally indistinguishable from segregating wild-types at all stages of development. Leaf initiation was unperturbed (Figure 1B, C), there was no loss of apical dominance or delayed flowering (Figure 1D, E), floral organs developed normally and the plants were fertile (Figure 1F, G). Furthermore, cross sections revealed no differences in patterning of the stem vasculature (Figure 1H, I) and paradermal views revealed normal leaf venation networks (Figure 1J-M). As such, contrary to previous reports, these results indicated that *AtDOT5* function is not necessary for leaf venation patterning or for the regulation of overall plant morphology.

### *The* Atdot5-2 *allele conditions no obvious developmental defects*

To try and reconcile our finding that AtDOT5 is dispensable for normal plant development with earlier reports, we re-evaluated the genotype and phenotype of the original *Atdot5-2* mutant line. The *Atdot5-2* allele was acquired as a homozygous line in the Col-0 background (SALK_148869c). In this line a T-DNA is inserted in the first exon of *AtDOT5* (At1g13290), upstream of the WIP and C2H2 zinc finger domains (Figure 2A). The genotype was confirmed as homozygous by genomic PCR using primers flanking the T-DNA insertion point together with a primer in the T-DNA left border (Figure S1). Importantly, the SALK_148869c line differs from the original SALK_148869 line in that the second insertion at the At2g26740 locus has been segregated away (Figure S1) and the mutant phenotype is not embryo lethal. The phenotype of the SALK_148869c mutant should thus reflect loss of function of *AtDOT5*. In experiments analogous to those performed with the gene edited *Atdot5* loss of function alleles (Figure 1), the *Atdot5-2* mutant phenotype was compared to that of Col-0 wild-type plants. *Atdot5-2* mutants exhibited normal leaf, rosette and flower development, no loss of apical dominance and full fertility (Figure 2B-G). Crucially, no changes were observed in either stem (Figure 2 H, I) or leaf vasculature patterning (Figure 2J-M). These data are consistent with our hypothesis that loss of *AtDOT5* function does not lead to perturbed vein patterning and suggest that the reported embryo lethal phenotype in the original SALK_148869 line (Petricka *et al*., 2008) was caused by loss of function of the soluble epoxide hydrolase encoded by the At2g26740 locus.

**Figure 2.**
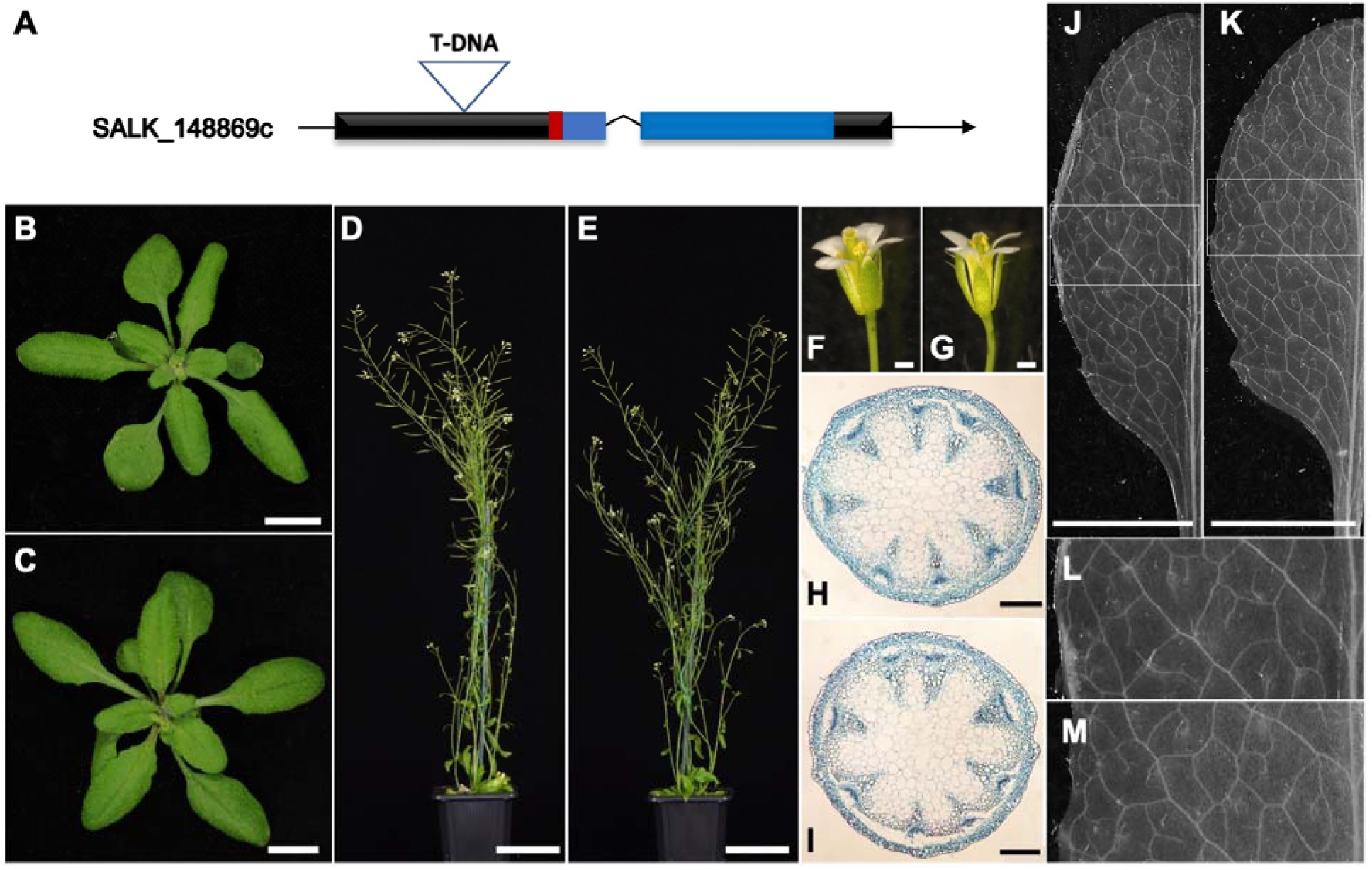
Homozygous *Atdot5-2* mutants (SALK_148869c) exhibit normal leaf venation patterns. **A)** Schematic of the *AtDOT5* locus showing the position of the T-DNA insertion in SALK_148869c line. Black boxes depict exons with the overlying red box depicting the WIP domain and the blue boxes depicting the zinc finger domains. **B-M)** Phenotypic characterization of Col-0 (B, D, F, H, J & L) and *Atdot5-2* mutants (C, E, G, I, K & M). Plants were imaged prior to bolting (B, C) and after flowering (D, E). Flower morphology appeared normal (F, G) and vascular patterning in both stems (H, I) and leaves (J-M) was similar in Col-0 and the T-DNA line SALK_148869c. L & M are insets of J & K respectively, as indicated. Scale bars: 1 cm (B, C); 5 cm (D, E); 0.5 mm (F, G); 200um (H, I) and 5 mm (J, K).

### *Developmental defects exhibited by the* Atdot5-1 *mutant cannot be explained by mutations in the* AtDOT5 *gene*

Given that the new gene edited alleles and the *Atdot5-2* allele, all of which are in the Col-0 background, do not lead to perturbed leaf venation patterns or to any general morphological defects, we considered whether the *Atdot5-1* mutant phenotype was specific to the L*er* background and thus to any genetic interactions present therein. To this end, we re-evaluated the phenotype and genotype of the original *Atdot5-1* mutant line.

Throughout development, *Atdot5-1* mutants exhibited a pleiotropic phenotype characterized by variable seedling morphology, altered phyllotaxy, delayed leaf initiation (Figure 3A, B), delayed flowering and loss of apical dominance (Figure 3C, D). The mutant also showed various defects in flower morphology that impacted on fertility and seed set (Figure 3E-G). Stamens either matured too quickly (Figure 3F) or did not elongate sufficiently (Figure 3G), failing in both cases to efficiently deliver viable pollen to the stigma. Additional vascular bundles were evident in the stem vasculature (Figure 3H, I) and the nearly glabrous and irregularly shaped leaves (Figure 3B) displayed conspicuous vein patterning defects (Figure 3J-M). Specifically, the leaf venation pattern was less complex in *Atdot5-1* leaves (Figure 3K, M) compared to wild type (Figure 3J, L) with fewer tertiary veins evident, most of the quaternary veins absent and higher order veins completely absent. These observations are consistent with the report that suggested *AtDOT5* influences multiple aspects of plant development, including leaf venation patterning (Petricka *et al*., 2008).

**Figure 3.**
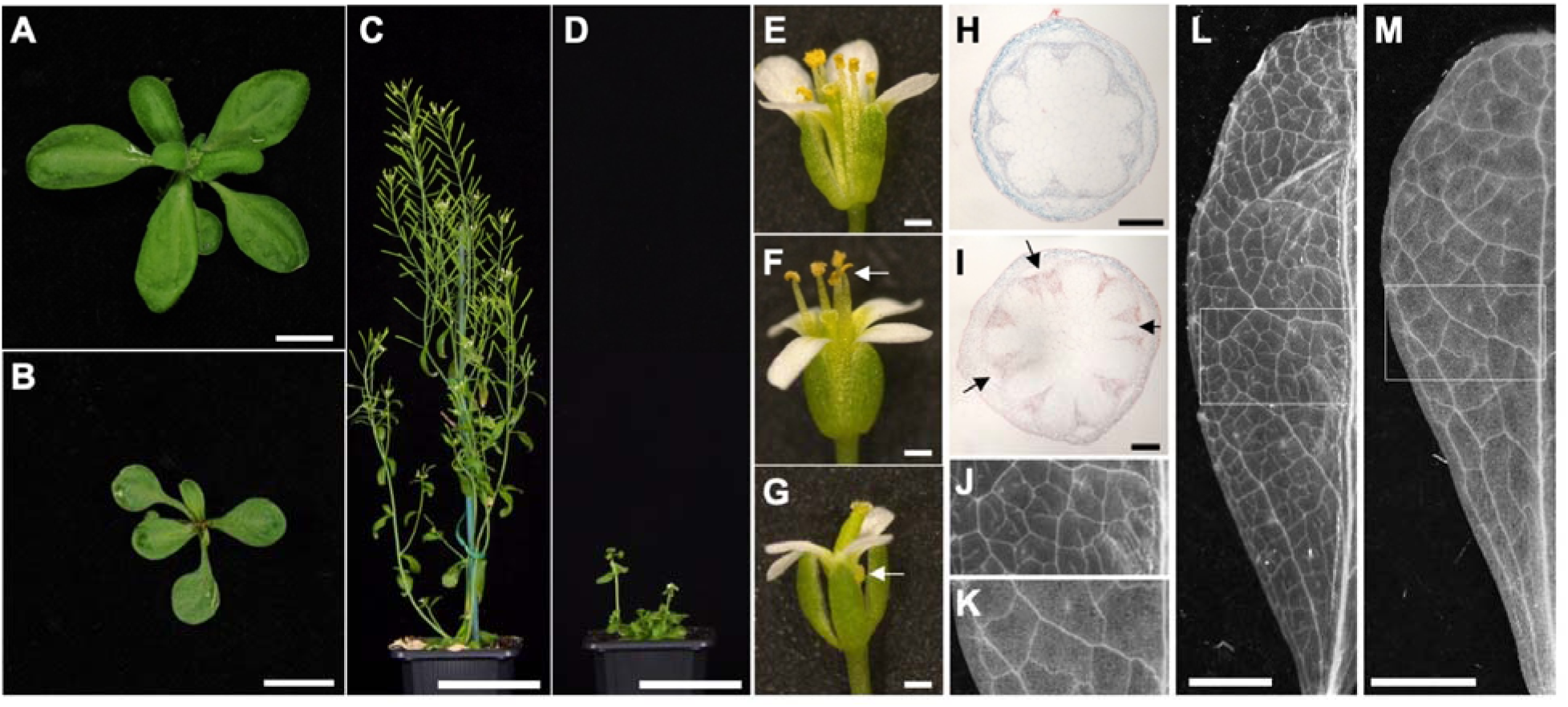
Pleiotropic phenotype of the *Atdot5-1* mutant. **A, B)** Contrasting rosette phenotypes of L*er* (A) and the *Atdot5-1* mutant (B) prior to flowering. **C-G)** Flowering phenotype of L*er* (C, E) and *Atdot5-1* mutant (D, F, G) plants. Mutant plants flower late and display reduced apical dominance (D). Floral morphology is perturbed with stamens either extending above the carpel (F - white arrow) or failing to fully emerge (G – white arrow). (**H-M)** Transverse sections through stems (H, I) and paradermal views of cleared juvenile leaves (J-M) from L*er* (H, J, L) and *Atdot5-1* mutant (I, K, M) plants. Additional vascular bundles are present in mutant stems (I - black arrows). J & K are insets of L & M respectively, as indicated. Scale bars 1 cm (A, B), 5 cm (C, D), 0.5 mm (E, F, G), 200μm (H, I) and 2.5 mm (L, M)

In an attempt to explain the conflicting phenotypic differences between *Atdot5-1* versus the gene edited and *Atdot5-2* loss of function mutants, we compared the *AtDOT5* locus in L*er* and Col-0 Arabidopsis accessions. To this end, the coding sequence of *AtDOT5* (At1g13750) from the recently published L*er* genome (Zapata *et al*., 2016) was aligned to the Col-0 *AtDOT5* (At1g13290) reference sequence (TAIR). The two sequences differ by 10 single nucleotide polymorphisms, six of which result in amino acid changes at positions 36 (T to A); 64 (S to T); 244 (S to G); 267 (V to E); 269 (E to K) and 294 (Y to C). In the original report by Petricka et al. (2008), nine point mutations were reported to alter the sequence of the *AtDOT5* gene in the *Atdot5-1* mutant background, with four of them resulting in amino acid changes at positions: 46, (T to A); 64 (S to T); 244 (S to G) and 294 (Y to C). The L*er* genome sequence reveals that three of the four amino acid changes reported as mutagenesis-induced in the *Atdot5-1* background correspond to natural variation present between L*er* and Col-0. We could not validate the fourth change (46 - T to A) in the genome of L*er* or in the genome of the *Atdot5-1* mutant (see below). As such, mutations in *AtDOT5* cannot explain the phenotypic changes observed in the *Atdot5-1* mutant.

### *Genome sequencing of the* Atdot5-1 *mutant reveals multiple polymorphisms and a Ds transposon insertion*

To identify potential causative mutations underlying the *Atdot5-1* phenotype, the genome of mutant plants was sequenced at 30-fold coverage. When sequence of the *AtDOT5* locus in the mutant background was aligned with the published L*er* genome, no significant changes were identified in *AtDOT5* or in any of the ten adjacent genes either upstream or downstream (Figure S2). As such, we cannot explain how partial complementation of the *Atdot5-1* mutant phenotype was achieved when sequences from this genomic region were used in transgenic complementation experiments (Petricka *et al*., 2008)

Given that the *Atdot5-1* line was identified in a collection of transposon-tagged mutants, we next looked for evidence of transgenes and/or transposons in the *Atdot5-1* mutant genome. A single transgene insertion was identified, corresponding to the transposon-containing construct used in the mutagenesis process (Bancroft *et al*., 1992). Transgene re-assembly from sequence traces showed that the *Ds* element was inserted in the coding sequence of the kanamycin resistance gene (*NPTII*) in the *Atdot5-1* mutant line, instead of in the streptomycin phosphotransferase gene where it was positioned in the original transformation construct (Figure 4A, B). This observation suggests that the *Ds* was transactivated by an autonomous *Activator* element at some point since the original lines were generated. Transposition of *Ds* is further evidenced by duplication of an 8 bp sequence ‘GCAGCTGT’ at the insertion point in *NPTII*. Although not well supported, a single pair of reads, one read corresponding to the *Ds* element and its paired read mapping to positions 11454793-11454942 on chromosome 4, suggests that a *Ds* element might also be inserted upstream of the final exon of ATLER-4G47010, the ortholog of AT4G20370 that encodes *TWIN SISTER OF FT (TSF*). However, because loss of function mutations in *TSF* do not condition phenotypes similar to *Atdot5-1* (Yamaguchi *et al*., 2005), and there was just a single read, we discounted the possibility that the insertion was significant. No other *Ds* insertion events were detected in the genome.

**Figure 4.**
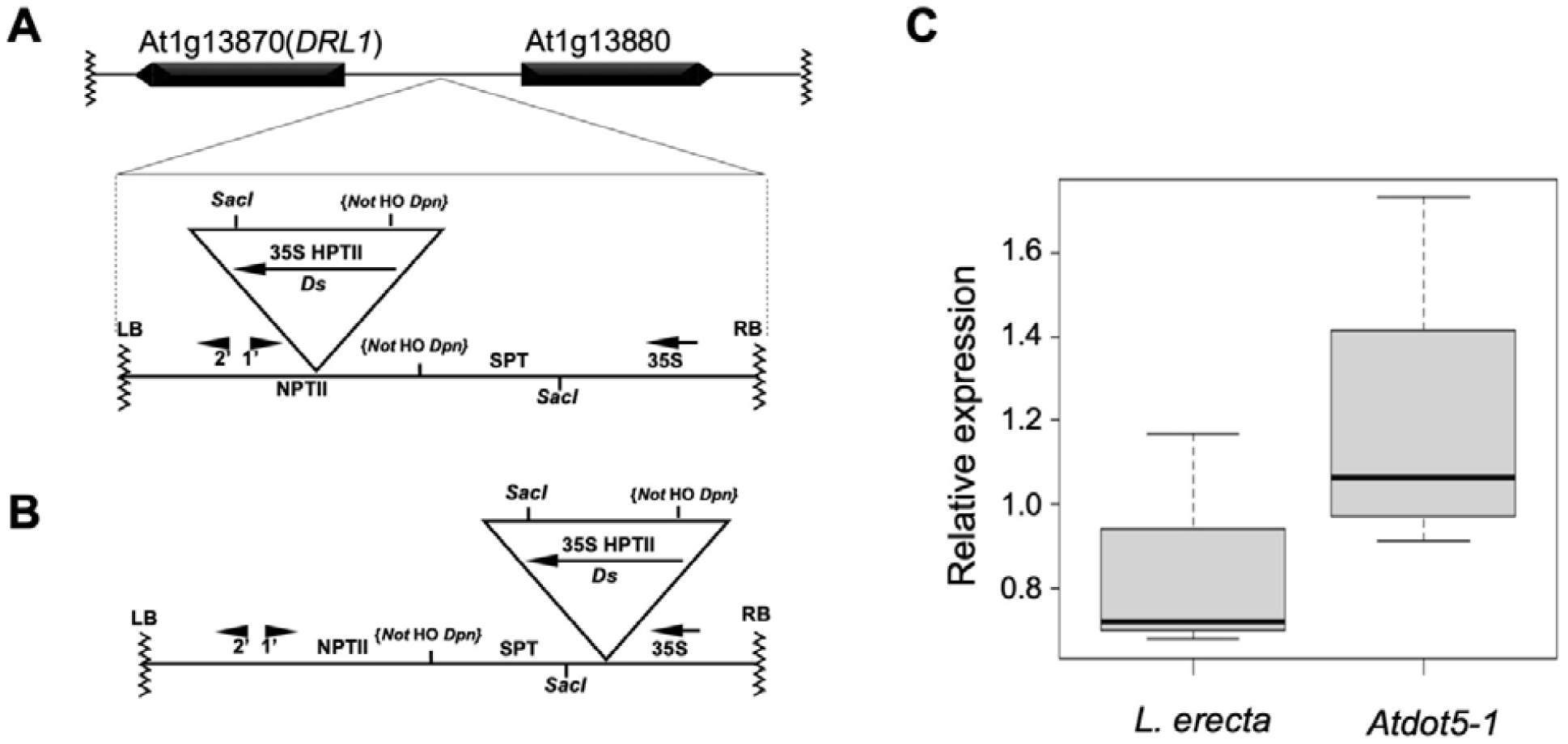
Transgene insertion on chromosome 1 in the *Atdot5-1* background. **A)** Diagram showing the insertion point of the Hm^R^ *Ds* construct in the *Atdot5-1* mutant genome, illustrating the position of the *Ds* in the *NPTII* gene. **B)** Diagram showing the original transformation construct as depicted by Bancroft et al. (1992) with the *Ds* inserted in the *SPT* gene. **C)** *DRL1* transcript levels in wild-type L*er* and *Atdot5-1* seedlings, as determined by qRT-PCR. Transcript levels were normalized against *Act2* transcript levels and an unpaired t-test demonstrated that there is no statistically significant difference between the two samples (P = 0.2395).

Sequence flanking the *Ds*-containing transgene in the *Atdot5-1* genome indicates that the transgene is inserted between the *DEFORMED ROOTS AND LEAVES (DRL1)* locus (At13870) and a gene of unknown function (At1G13880) (Figure 4A). *DRL1* was first identified by *Ds* tagging with the reported loss of function *drl1-1* mutant caused by a transposed *Ds* (*tDs*) in the *DRL1* coding sequence (Bancroft *et al*., 1993). Genomic PCR confirmed that the *Ds*-containing transgene is present at the same genomic location in both *Atdot5-1* and *drl1-1* mutants, suggesting that it may have been the source of the *tDs* that inserted into the *DRL1* coding region (Figure S3). Although some aspects of the *Atdot5-1* mutant phenotype resemble those found in *drl1* mutants, some are noticeably different. For example, *drl1-1* mutants do not form inflorescences (Bancroft *et al*., 1993) and *drl1-2* mutants show no venation patterning defects even though leaves are narrower than wild-type (Nelissen *et al*., 2003). As such, we hypothesized that the transgene insertion upstream of the *DRL1* locus in *Atdot5-1* has no functional significance. This hypothesis is supported by the fact that any phenotypic consequences of the insertion would have been segregating in the progenitor lines of the *drl1-1* mutant, and none were reported (Bancroft *et al*., 1993). As a final verification, we assessed whether *DRL1* gene expression is perturbed in *Atdot5-1* mutants. *DRL1* encodes a putative elongator associated protein that is expressed in all organs during wild-type development (Nelissen *et al*., 2003). Notably, qRT-PCR using RNA extracted from pooled 7d old seedlings demonstrated that transcript levels are not significantly different between wild-type L*er* and *Atdot5-1* mutant lines (Figure 4C), and the genome sequence reveals no mutations in the *DRL1* coding sequence. We thus conclude that the pleiotropic *Atdot5-1* mutant phenotype is not caused by a transposon or transgene insertion in the genome, nor by any loss or gain of *DRL1* function

To identify potential causative mutations in the *Atdot5-1* genome, sequence variations that have the potential to disrupt gene function were identified. A total of 8 exonic frameshift deletions and 10 exonic frameshift insertions were identified (Table 1). Previous reports of phenotypes associated with loss of function at two of the identified loci - AT3G49360 (Xiong *et al*., 2009) and At5G54600 (Liu *et al*., 2013) – suggest that the frameshifts observed at these loci are not responsible for the *Atdot5-1* phenotype. The other sequence variants cannot be eliminated as potential causative mutations without further investigation.

**Table 1.** Frameshift mutations identified in the *Atdot5-1* genome.

| ANNOTATION | TYPE of MUTATION |  | CHR | TAIR 10 | DESCRIPTION |
| --- | --- | --- | --- | --- | --- |
| AtLer-1G22960.1 | deletion | before ATG | chr1 | AT1G12330 | cyclin-dependent kinase-like protein |
| AtLer-1G22970.1 | deletion | exon3 | chr1 | not annotated |  |
| AtLer-1G22970.1 | deletion | exon4 | chr1 |  |  |
| AtLer-1G38900.1 | deletion | 11bp deletion, heterozygous | chr1 | AT1G27580 | F-box protein (DUF295) |
| AtLer-3G50050.1 | deletion | exon7 | chr3 | not annotated |  |
| AtLer-4G42200.1 | deletion | exon | chr4 | AT4G16230 | GDSL-like Lipase/Acylhydrolase superfamily protein |
| AtLer-4G62980.1 | deletion | exon 4, premature STOP | chr4 | AT4g34200 | PGDH1; phosphoglycerate dehydrogenase 1; EDA9 |
| AtLer-5G70960.1 | deletion |  | chr5 | AT5G56200 | C2H2 type zinc finger transcription factor family |
| AtLer-1G22980.1 | insertion | exon 2 | chr1 | not annotated |  |
| AtLer-1G47950.1 | insertion |  | chr1 | not annotated |  |
| AtLer-1G78630.1 | insertion | before ATG | chr1 | AT1G65295 |  |
| AtLer-2G35290.1 | insertion | no STOP | chr2 | AT2G18160 | ATBZIP2; basic leucine-zipper 2; FLORAL TRANSITION AT THE MERISTEM3; FTM3; GBF5; G-BOX BINDING FACTOR 5; bZIP2 |
| AtLer-3G18110.1 | insertion | before ATG | chr3 | AT3G07970 | Pectin lyase-like superfamily protein |
| AtLer-3G27460.1 | insertion | before ATG | chr3 | AT3G16340 | Taurine-transporting ATPase |
| AtLer-3G63470.1 | insertion | exon 4 | chr3 | not annotated |  |
| AtLer-3G68040.1 | insertion | premature stop | chr3 | AT3G49360 | PGL2. Acts redundantly with PGL1 and PGL5 (Xiong et al., 2009) |
| AtLer-5G23930.1 | insertion | before ATG | chr5 | AT5G14200 | isopropylmalate dehydrogenase 1 |
| AtLer-5G74960.1 | insertion |  | chr5 | AT5G54600 | PLASTID RIBOSOMAL PROTEIN L24, RPL24, SUPPRESSOR OF VARIEGATION 8, SVR8 (Liu et al., 2013) |

## CONCLUSION

Gene editing technologies allow for unambiguous assignment of mutant phenotypes to loss of function alleles. This advance has enabled more robust hypotheses of gene function to be proposed than was previously possible. Genetic screens that utilized highly mutagenic chemicals or non-specific insertion tags such as T-DNA or transposons inevitably led to ‘noisy’ genomes that could mask the actual mutation of interest, and RNAi suppression lines similarly led to imprecise interpretations of gene function. The discovery by gene editing that the long-standing AUXIN BINDING PROTEIN played no role in auxin homeostasis is probably one of the most high-profile cases of mistaken identity in plant biology (Gao *et al*., 2015) but there will undoubtedly be more over the coming years. In this context, we have shown here that the *AtDOT5* gene does not regulate venation patterning in the Arabidopsis leaf.

## EXPERIMENTAL PROCEDURES

### Plant material

*Atdot5-1* seeds were obtained from David Diaz Ramirez and Nayelli Marsch Martinez (Biotecnology and Biochemistry Department, Centre for Research and Advanced Studies (CINVESTAV-IPN) Irapuato Unit, Mexico). The homozygous T-DNA line SALK_148869c was obtained from NASC (stock number N648869). Landsberg (L*er*) and Columbia_0 (Col-0) seeds were originally obtained from Lehle Seeds but have been propagated in the lab for many years. Seeds were sown directly on soil (Levington Seed modular compost), stratified at 4°C for two days to break dormancy and then transferred to a controlled environment chamber (CER) with a set temperature of 21°C and a 16-hour photoperiod. Col-0 plants used to generate CRISPR lines were grown in the greenhouse under the same temperature and photoperiod conditions as the CER.

### Construct design and plant transformation

A short RNA guide (sgRNA) was designed against the first exon of At1g13290 gene reference sequence using the CRISPOR online tool (Concordet and Haeussler, 2018). Constructs were generated using parts of a modular cloning system based on Golden Gate technology (Weber *et al*., 2011). The guide sequence was integrated by PCR into an RNA scaffold derived from EC15768 (Hughes and Langdale, 2022), modified by the addition of 34 bp 5’-CTAGACCCAGCTTTCTTGTACAAAGTTGGCATTA-3’ at the 3’ end as found on pICH86966 (Nekrasov *et al*., 2013). The scaffold was assembled into a Golden Gate level 1 module (position 3, reverse) downstream of the AtU6-26 promoter. The promoter was amplified by PCR from Col-0 genomic DNA using primers: AtU6-26pF: 5’-cactctgtggtctca<u>GGAG</u>AAGCTTCGTTGAACAACGGA-3’ and AtU6-26pR 5’-cactctgtggtctca<u>CAAT</u>CACTACTTCGACTCTAGCTG-3’ containing 4 bp sequences and *BpiI* restriction sites compatible with the PU Level 0 vector, EC41295.

The Cas9p gene (Ma *et al*., 2015) obtained by synthesis as an SC module (CDS) was cloned under the control of the AtYAO (At4g05410) promoter (Yan *et al*., 2015) in a Golden Gate level 1 module corresponding to position 2, forward. The AtYAO promoter was obtained by PCR from genomic DNA with primers pAtYAO-F 5’-cactctgtggtctca<u>GGAG</u>ACCCAAATCAACAGCTGCAA-3’ and pAtYAO-R 5’-cactctgtggtctcaC<u>ATTT</u>CTTCTCTCTCTCACTCCCTCT-3’ and cloned as described above into a PU Level 0 vector, EC41295. To terminate transcription the t-NOS terminator module, EC41421, was used. To allow for the selection of transgenic plants, a module containing the bar gene fused to the *A. tumefaciens* nopaline synthase (NOS) promoter and terminator was cloned adjacent to the LB of the pICSL4723 Golden Gate level 2 backbone, in position 1, reverse. Arabidopsis Col-0 plants were transformed by floral dipping (Clough and Bent, 1998).

### Genotyping

To genotype CRISPR lines, 96 T1 basta resistant seedlings were analysed using a CAPS marker designed to cut the wild type sequence at the predicted editing site (3 bp into the guide sequence from the PAM) (Figure S4). The undigested fragment was sequenced to identify the nature of the mutations. The T2 progeny were screened to select mutant lines free of the transgene. When it was not possible to segregate the mutation from the transgene, the lines were backcrossed onto Col-0.

### Genome sequence

DNA was isolated from pooled 3 week old *Atdot5-1* mutant seedlings and sequenced. Plant WGS was performed by Novogene Cambridge using a standard Illumina pair-end (PE) sequencing protocol with 30X coverage. 5G data was obtained and mapped to the *Landsberg erecta* genome available from the 1001 Genomes Browser: https://1001genomes.org/data/MPIPZ/MPIPZJiao2020/releases/current/strains/Ler/. Standard data analysis provided by Novogene included extracting polymorphisms and predicting their impact on gene function. Lists of single nucleotide polymorphisms (SNP), insertions/deletions (InDel), structural variation (SV) and copy number variation analysis (CNV) were included in the results. Limitations on the bioinformatics pipeline did not allow mapping of the transgene and transposon insertions which were assembled and mapped by manually searching sequence traces.

### qRT-PCR

Total RNA was extracted from pooled 7-day old seedlings using the Qiagen RNeasy Plant Mini Kit and complementary DNA was synthesized using the Maxima cDNA Synthesis kit (Thermofisher). Quantitative RT-PCR analysis was performed on a StepOnePlus Real-Time PCR System (Applied Biosystems) using the SYBR™ Green PCR Master Mix (Ref. 4309155, applied biosystems by Thermo Fisher Scientific) with cycle conditions 95°C for 10 min, then 40 cycles of 95°C for 15 s, 60°C for 10 s and 72 °C for 15 s.

To amplify the *DRL1* gene, primers were designed against the CDS using Primer3Plus (DRL1qRT-F : 5’ – GTTGGACAGAGCGACACAAG-3’, DRL1qRT-F : 5’ – GTGGACCGCTTAGACTCGAT - 3’) and expression levels were normalized to the expression of *Actin2* amplified using previously published primers (Liu *et al*., 2009).

Three biological replicates for *L. erecta*, 4 biological replicates for *Atdot5-1* mutant and three technical replicates for each sample were run. The Cq values and primer efficiencies were calculated for each sample using the R package qpcR (Ritz and Spiess, 2008) and normalization and relative quantification of expression were calculated using the EasyqpcR package based on previously published algorithms (Hellemans *et al*., 2007). The box plot was generated using RStudio.

### Histology

To analyse leaf venation patterns, leaf 5 from ∼3 week old (CRISPR mutants, Salk lines, wild type L*er* and Col-0) or ∼4 week old (*Atdot5-1*) plants was fixed in 3:1 ethanol:acetic acid and chlorophyll was cleared by successive washes with 70% ethanol followed by overnight incubation in histoclear. Leaf vasculature was imaged using a Leica S9i stereomicroscope against a dark background.

Young stem segments were cut 1 cm from the base of the inflorescence soon after bolting. Where possible the plants were selected to have stems of equivalent heights. In the case of the *Atdot5-1* mutant and L*er* wild type control, lateral shoots were used instead of the main inflorescence because the mutant lacked apical dominance. One cm segments were fixed overnight using 3:1 ethanol:acetic acid and then infiltrated with paraffin wax in a Tissue-Tek VIP machine (Sakura, www.sakurag.eu) following a protocol described previously (Hughes *et al*., 2019). 10μm sections were stained using a 1% Safranin solution in 50% ethanol, counterstained with a 0.04% fast-green solution in 95% ethanol and mounted in DPX mounting medium. Brightfield images were obtained using the Leica LASX software and a DFCT000T camera fitted on a Leica DMRB microscope.

## AUTHOR CONTRIBUTIONS

DV and JAL conceived and designed the experiments. DV carried out the experiments and analysed the data. DV and JAL wrote the manuscript.

## ACKNOWLEDGEMENTS

We are grateful to Julie Bull, Roxaana Clayton and Lizzie Jamieson for technical support; John Baker for photography; Steve Kelly for help with bioinformatic analyses; and Tom Hughes, Sophie Johnson, Julia Lambret-Frotte, Chiara Perico, Sovanna Tan and Maricris Zaidem for discussion throughout the experimental work and during manuscript preparation. This work was funded by the Bill and Melinda Gates Foundation C_4_ Rice grant awarded to the University of Oxford (2015-2019, OPP1129902; 2019-2024, INV-002970).

## DATA STATEMENT

All data generated or analysed during this study are included in this published article and its supplementary information files, except for the sequence reads of the *Atdot5-1* mutant genome which are available at ArrayExpress https://www.ebi.ac.uk/arrayexpress - accession number E-MTAB-12010.

## SUPPORTING INFORMATION

**Figure S1**. Validation of T-DNA insertion in *AtDOT5* in SALK_148869c line.

**Figure S2**. Snapshots of sequencing reads for genes flanking *AtDOT5* in the *Atdot5-1* genome.

**Figure S3**. Confirmation of shared transgene position in *Atdot5-1* and *Atdrl1-1* mutants.

**Figure S4**. PCR assay to genotype gene edited alleles of *AtDOT5*.

## SUPPORTING INFORMATION

**Figure S1.**
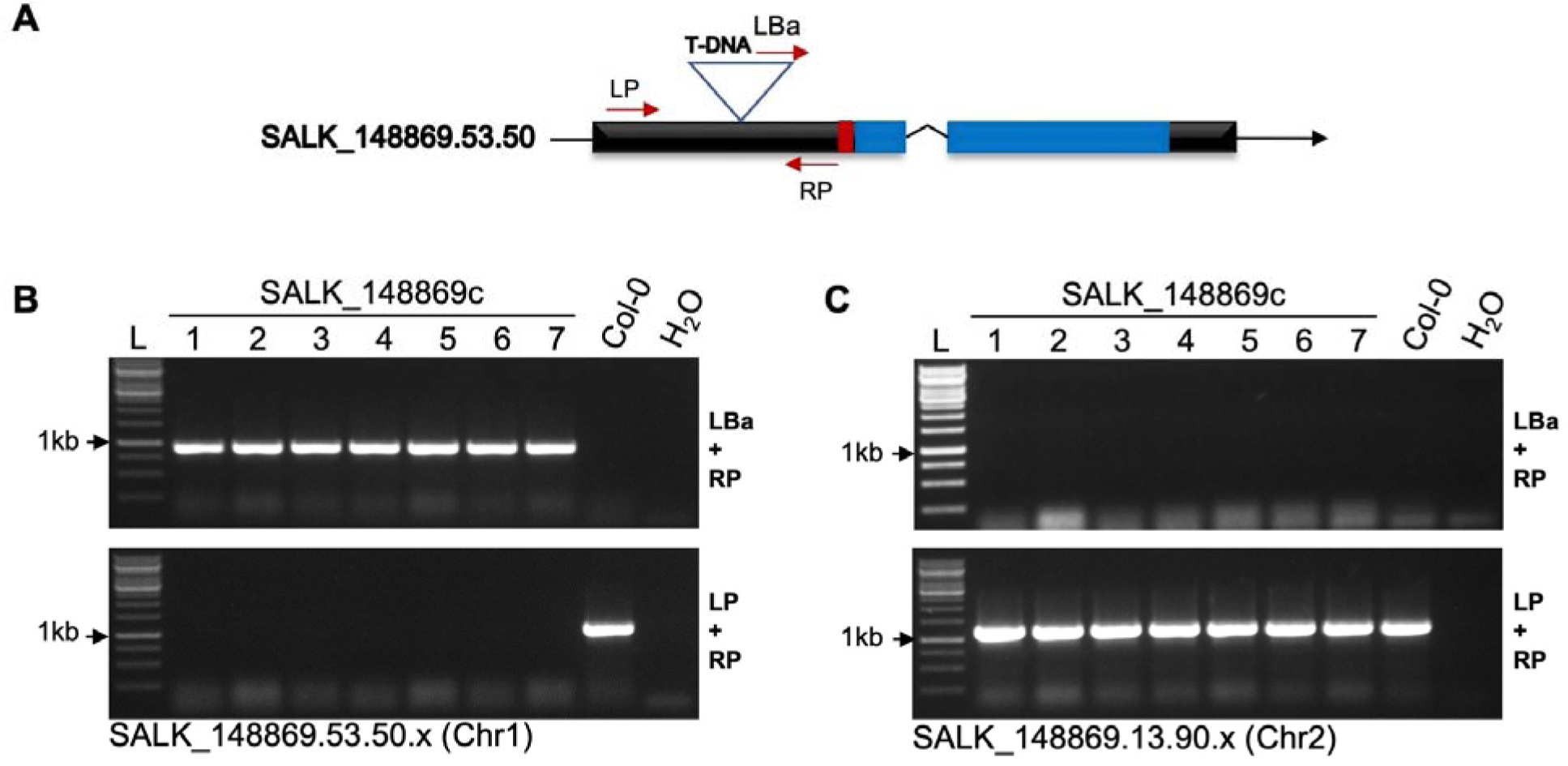
Validation of the T-DNA insertion at the *AtDOT5* locus in the homozygous SALK_148869c line. **A)** Schematic representation of the T-DNA insertion at the *AtDOT5* locus (SALK_148869.53.50.x). Black boxes depict exons with the overlying red box depicting the WIP domain and the blue boxes depicting the zinc finger domains. Inverted triangle marks the position of the T-DNA in the first exon and the red arrows show the positioning of the genotyping primers. **B)** PCR validation of T-DNA insertion at the *AtDOT5* locus (At1g13290). Primers were designed in the T-DNA border (LBa1: 5’-TGGTTCACGTAGTGGGCCATCG-3’) and in the first exon of *AtDOT5* both upstream (LP: 5’-TTCCCACAATTTTTCTCATGC-3’) and downstream (RP: 5’-CATGGGCTTGTCACTCGTAAC -3’) of the T-DNA insertion. The expected 438-738 bp fragment was amplified by PCR using LBa and RP primers. Amplification using the LP and RP primers yielded the expected 1062 bp fragment in Col-0 but no fragments in the SALK_148869c line due to the presence of the T-DNA insertion. The SALK_148869c line is thus homozygous for the T-DNA insertion at *AtDOT5*. **C)** PCR showing absence of a T-DNA insertion at the soluble epoxide hydrolase encoding At2g26740 locus. Because an additional insertion (SALK_148869.13.90.x) was reported in the parental line SALK_148869, primers were designed to amplify sequences from the At2g26740 locus (in an analogous design to that for *AtDOT5*) in the SALK_148869c line. No fragments were amplified using the LBa1 and RP (5’-TCAATTTGGTTAATGATTTGCC-3’) primers, demonstrating the absence of a T-DNA insertion at the locus. By contrast, a 1172 bp fragment was amplified from all SALK-148869c individuals and from Col-0 using RP and LP (5’-CCCCAAAAACTTTTGTCCTTC-3’) primers.

**Figure S2.**
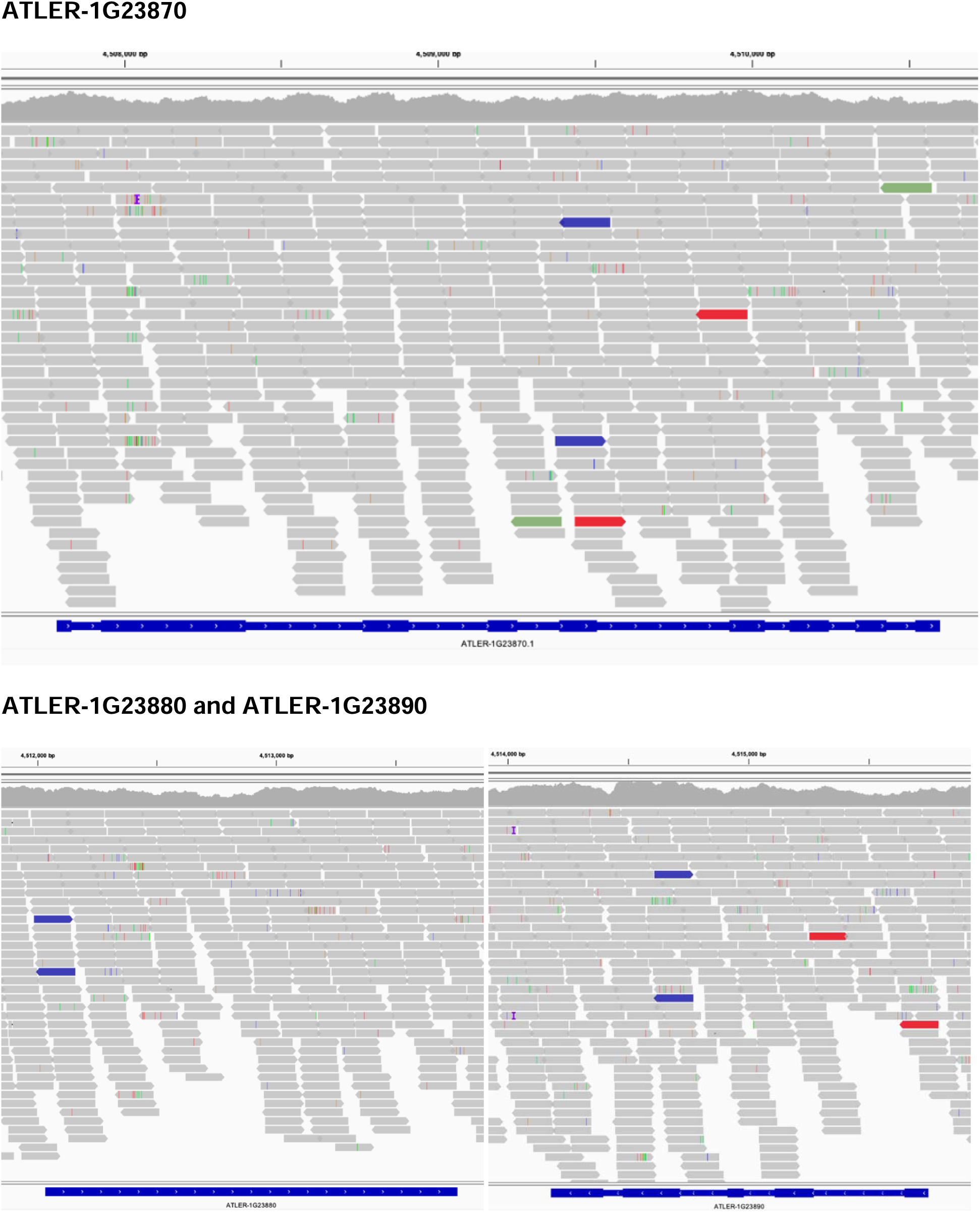
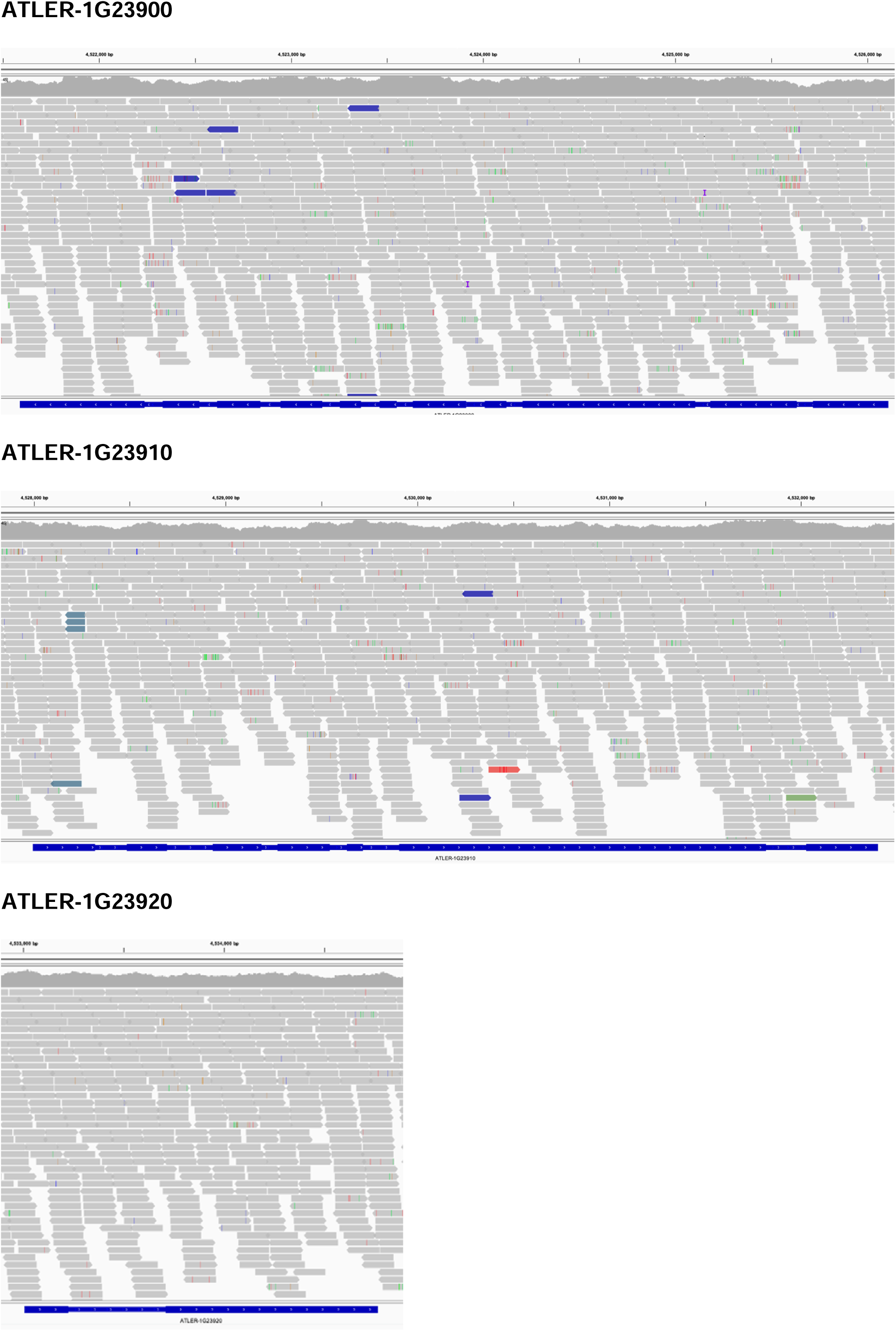
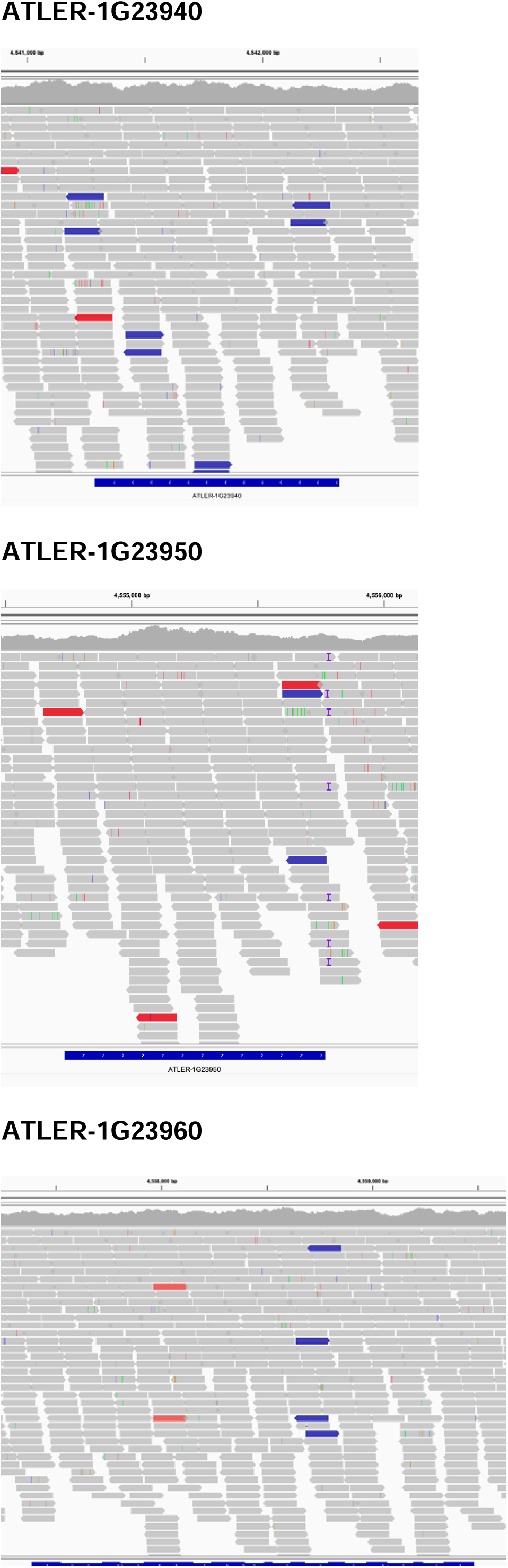
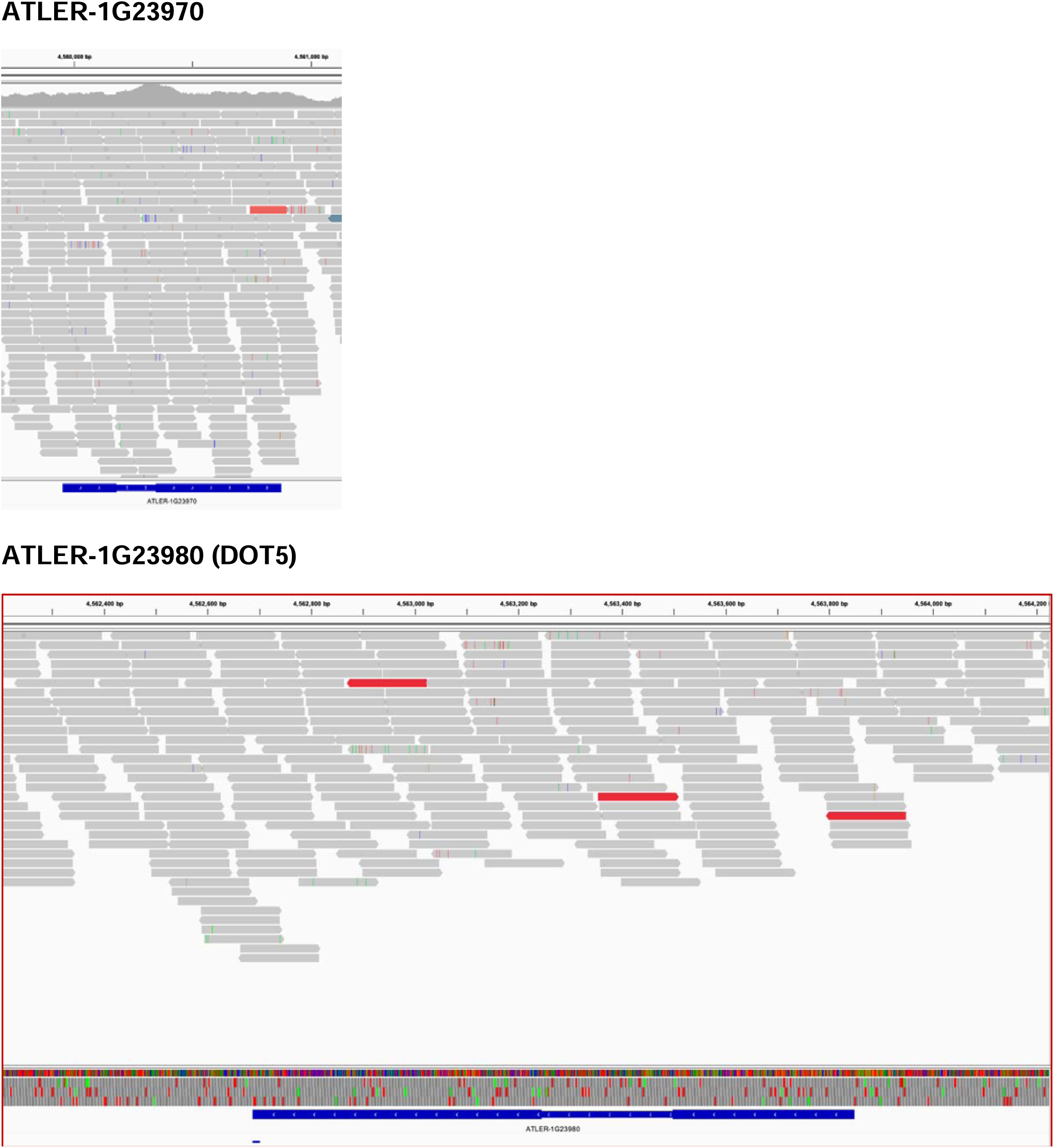
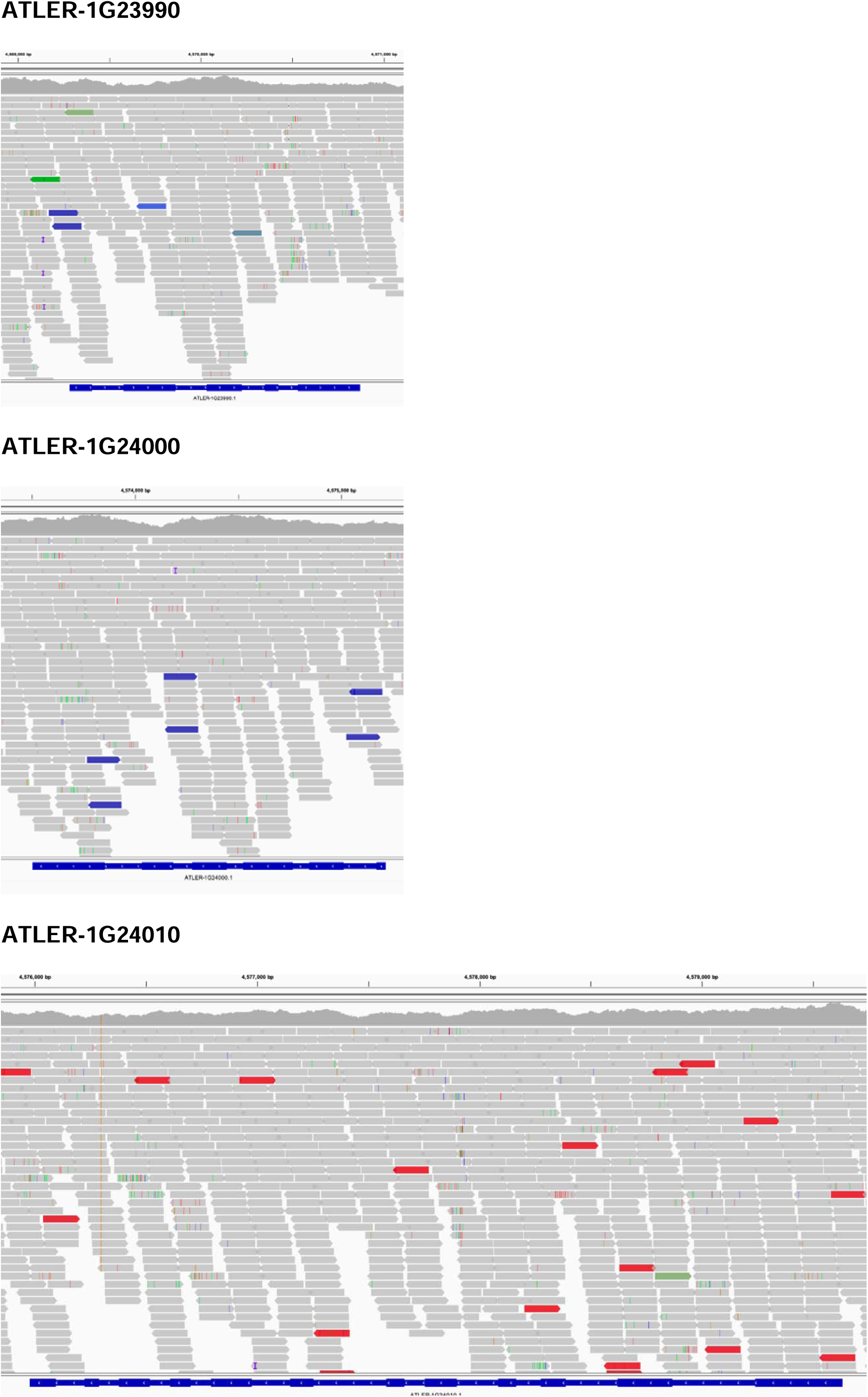
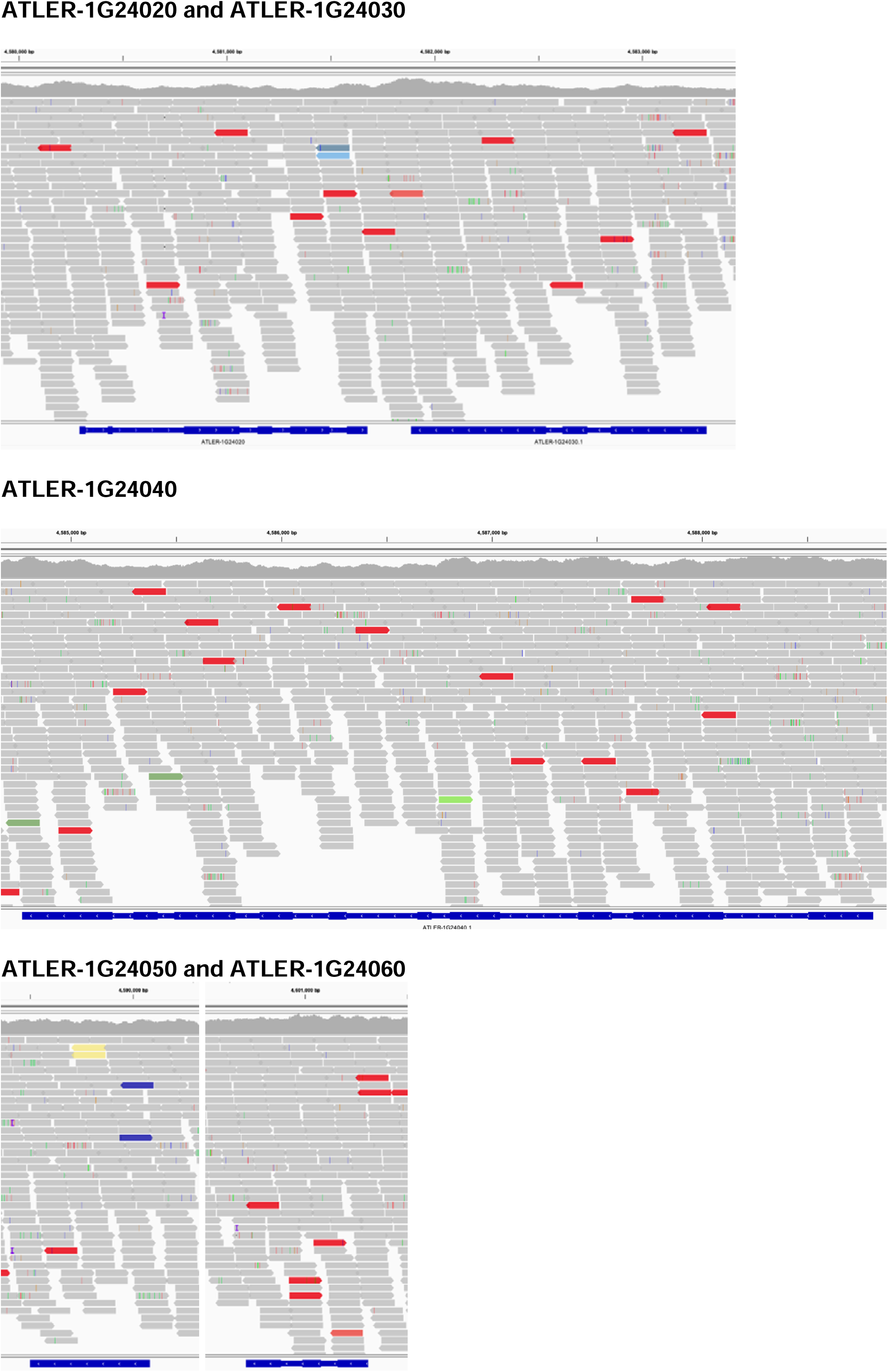
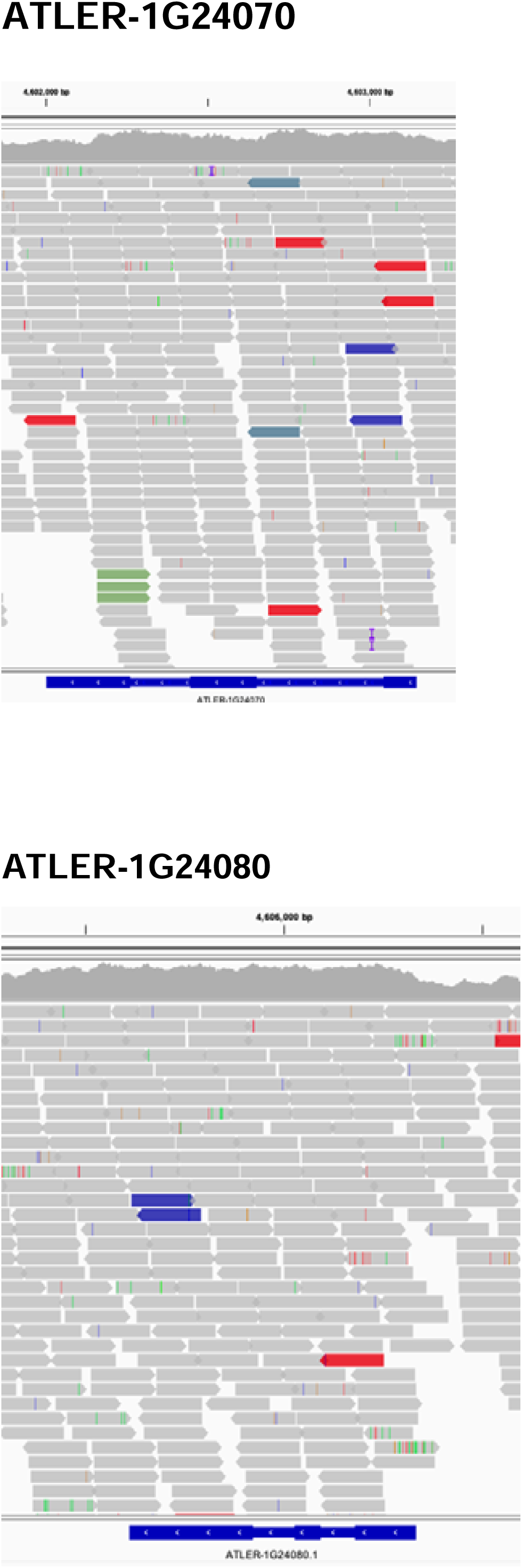
Snapshots of the sequencing reads for ten genes flanking either side of *AtDOT5*, aligned to the Landsberg *erecta* reference genome. Reads were visualized using the *Integrative Genomics Viewer (IGV)* software. Reads are shown as arrows, the blue bars at the bottom of the image are gene models with wider regions representing exons.

**Figure S3.**
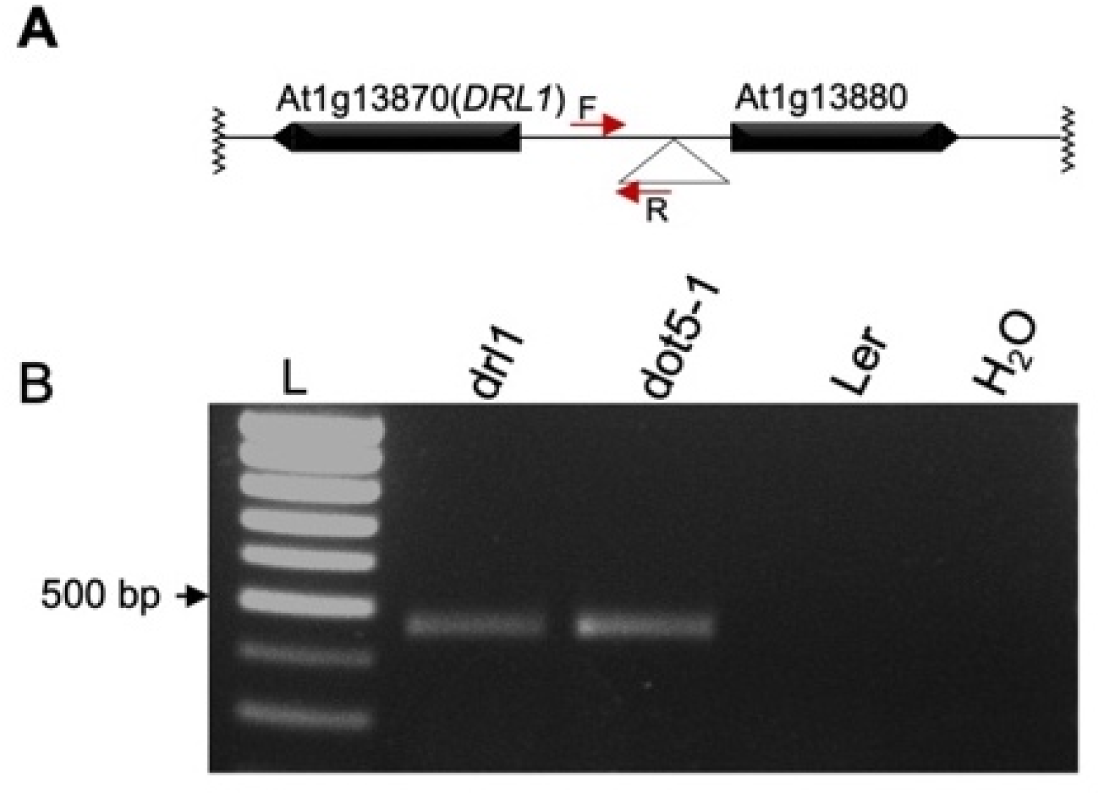
Confirmation that the *Ds-*containing transgene insertion is in the same position in *Atdrl1-1* and *Atdot5-1* mutants. **A)** Schematic representation of the *Ds* insertion in the *DRL1* promoter region. Inverted triangle marks the position of the insertion and the red arrows show the position of the forward (F) and reverse (R) genotyping primers used for PCR. **B)** PCR amplicons of similar sizes were obtained from both the *Atdrl1-1* and *Atdot5-1* mutants suggesting that the insertion is present at the same location in both mutant backgrounds.

**Figure S4.**
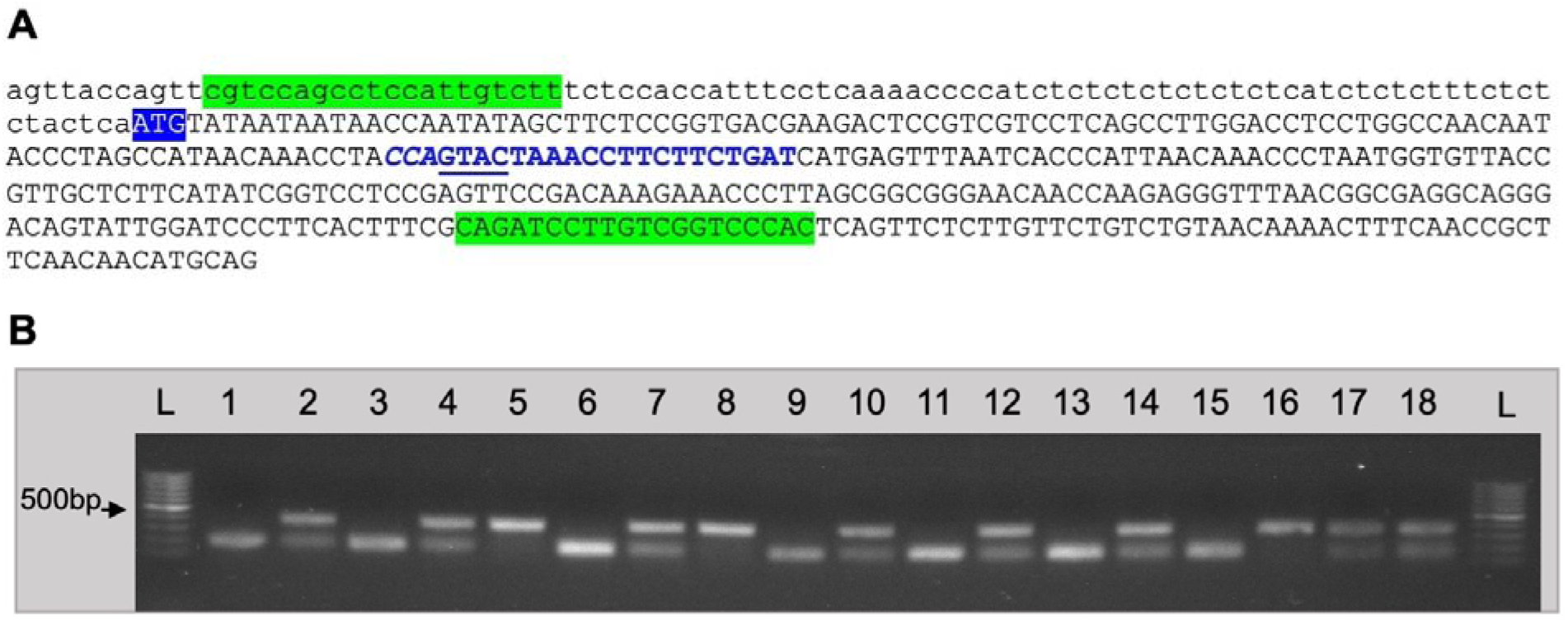
Genotyping of the *AtDOT5* CRISPR alleles. **A)** Sequence containing the first exon showing the position of the guide sequence (reverse) highlighted in blue with the PAM site shown in italics and the restriction site for *RsaI* (GTAC) underlined. Cas9 will induce mutations next to the PAM. If a mutation is induced, *RsaI* is unlikely to cut the PCR product amplified from the mutant sequence. Primers are highlighted in green. **B)** Example of an *RsaI* digest of fragments amplified from a segregating T3 population (samples 1 to 18). The lower fully digested band is amplified from the wild type allele and the upper undigested band from the mutant allele. L indicates the molecular weight marker.

